# Local expectation violations result in global activity gain in primary visual cortex

**DOI:** 10.1101/067579

**Authors:** Peter Kok, Lieke L.F. van Lieshout, Floris P. de Lange

## Abstract

During natural perception, we often form expectations about upcoming input. These expectations are usually multifaceted – we expect a particular object at a particular location. However, expectations about spatial location and stimulus features have mostly been studied in isolation, and it is unclear whether feature-based expectation can be spatially specific. Interestingly, feature-based attention automatically spreads to unattended locations. It is still an open question whether the neural mechanisms underlying feature-based expectation differ from those underlying feature-based attention. Therefore, establishing whether the effects of feature-based expectation are spatially specific may inform this debate. Here, we investigated this by inducing expectations of a specific stimulus feature at a specific location, and probing the effects on sensory processing across the visual field using fMRI. We found an enhanced sensory response for unexpected stimuli, which was elicited only when there was a violation of expectation at the specific location where participants formed a stimulus expectation. The neural consequences of this expectation violation, however, spread to cortical locations processing the stimulus in the opposite hemifield. This suggests that an expectation violation at one location in the visual world can lead to a spatially non-specific gain increase across the visual field.

## Introduction

As you hear the sound of the ice cream van approaching around the corner, you have a strong expectation of *what* you will see next, as well as *where* you will see it. Such spatially as well as feature specific expectations are ubiquitous in natural perception. However, expectations about spatial location and stimulus features have mostly been studied in isolation. Independently of each other, both spatial^1,2^ and feature-based^3,4^ expectation have been shown to affect neural processing in early sensory regions. Generally, a suppression of activity is observed when expectations are met by congruent input, and an increase in activity is observed when expectations are invalid^1^. However, it is unclear whether the neural modulation by feature-based expectation (e.g., expecting the colour red) can be spatially specific. This question is particularly interesting given that the higher-order regions in frontal cortex containing spatial and feature information may be largely separate^5^. It is currently unknown whether these regions operate independently, or whether they can synergistically induce top-down modulations in early visual cortex that are both spatially and feature specific^6^. That is, whether feature-based expectation can affect processing of stimuli at the location in which the stimulus is expected to occur, without affecting processing of stimuli presented elsewhere in the visual field.

Of note, feature-based attention has been shown to have spatially non-specific effects, spreading across the visual field^7–11^, affecting even the processing of fully irrelevant peripheral stimuli^12^. However, feature-based attention and expectation have been argued to be dissociable^13^, and their neural mechanisms may be distinct^3,14^. Establishing whether or not the modulatory effects of feature-based expectation on sensory responses, unlike those of feature-based attention, are spatially specific may help to shed light on this much debated issue.

In the current study, we investigated the spatial specificity of feature-based expectation by inducing expectations of a specific stimulus feature (i.e., orientation) at a specific location, and probing the effects on neural sensory processing across the visual field. To preview, we found that expectation effects were only evoked by unexpected features presented at the expectation-specific location, and not by features presented at the opposite location. The *consequences* of an expectation violation, however, were found to be spatially non-specific, in that the increased neural response was present at both cortical locations processing stimuli.

## Materials and Methods

### Participants

Thirty healthy individuals participated in the experiment. All participants were righthanded, MR-compatible and had normal or corrected-to-normal vision. The study was approved by the local ethics committee (CMO Arnhem-Nijmegen, The Netherlands) under the general ethics approval (“Imaging Human Cognition”, CMO 2014/288), and the experiment was conducted in accordance with these guidelines. All participants gave written informed consent according to the declaration of Helsinki. One participant was excluded from analysis due to excessive (> 5 mm) head movements in the scanner and one participant was excluded due to the absence of a clear visual signal during the retinotopic mapping session, precluding the drawing of ROIs. Five participants were excluded due to failure to perform one or both of the tasks with above chance accuracy. The final sample consisted of 23 participants (15 female, age 23 ± 3, mean ± SD), who completed both a behavioural and an fMRI session.

### Stimuli

Visual stimuli were generated using MATLAB (Mathworks, Natick, MA, USA) and the Psychophysics Toolbox^15^. In the behavioural session, the stimuli were presented on a BENQ XL2420T monitor (1024 x 768 resolution, 60 Hz refresh rate). In the fMRI session, the stimuli were displayed on a rear-projection screen using a luminance calibrated EIKI (EIKI, Rancho Santa Margarita, CA) LC-XL100 projector (1024 x 768 resolution, 60 Hz refresh rate) against a uniform grey background. During the fMRI session, participants viewed the visual display through a mirror that was mounted on the head coil. The visual stimuli consisted of two dots (one coloured, either orange or cyan, and one grey, calibrated to be of equal luminance) and two pairs of greyscale luminance-defined sinusoidal Gabor grating stimuli (Figure 1). The two dots (0.7° diameter) were centred respectively 1° left and 1° right of the fixation bull’s eye and the grating stimuli were centred at 5° of visual angle to the left and to the right of fixation (grating radius = 3.5°, contrast decreased linearly to zero over the outer 0.5°). At each grating location (i.e. left and right of fixation), the two consecutively presented gratings differed from each other in terms of phase, spatial frequency, orientation, and contrast. The first grating had random spatial phase, and the second grating was in anti-phase to the first grating. The two gratings had spatial frequencies of 1.0 and 1.5 cpd, respectively, the order of which was pseudorandomised and counterbalanced over conditions. The first grating had an orientation of either 45° or 135°, and the second grating was tilted slightly clockwise or anti-clockwise with respect to the first. The mean Michelson contrast of the two gratings was 80%, with one grating being slightly lower and the other slightly higher than that. Whether the first or second grating had the lower contrast was pseudorandomised and counterbalanced over conditions. The exact orientation and contrast differences between the two gratings was determined using an adaptive staircase procedure in the behavioural session, see below. The direction of the spatial frequency, contrast, and orientation changes were fully independent (counterbalanced over conditions) for the two grating locations.

**Figure 1.**
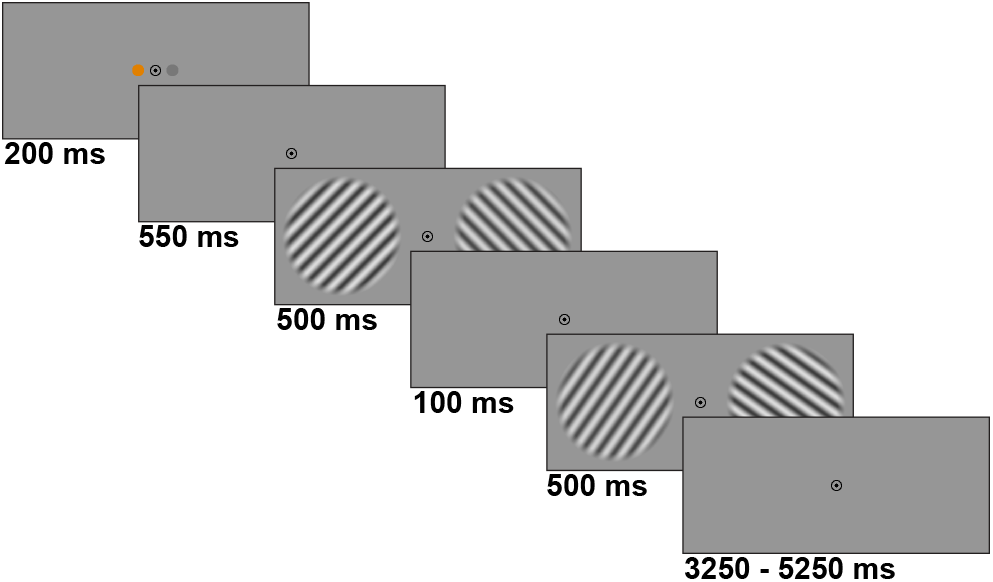
Experimental paradigm. The coloured dot indicated which side of the screen was relevant, and the specific colour (orange or cyan) predicted the orientation of the gratings on that side (75% valid). Participants performed either a contrast or orientation discrimination task on the two consecutive gratings on the attended side (ignoring the gratings presented on the other side), in separate scanner runs. Importantly, the colour of the cued dot bore no relationship with the orientation of the gratings presented at the non-cued side.

### Experimental design

Each trial of the main experiment started with the presentation of two dots (one grey and the other either orange or cyan) appearing next to the fixation bull’s eye for 200 ms. After a 550 ms delay, two pairs of consecutive grating stimuli were presented for 500 ms each, separated by a blank screen (100 ms) (see Figure 1). The intertrial interval (ITI) was jittered between 3250 and 5250 ms. The fixation bull’s eye was present throughout the entire trial. Participants were instructed to maintain fixation on the central bull’s eye during each experimental run and to covertly attend to one of the two grating locations. Participants were instructed to attend to the grating at the side of the coloured dot, and ignore the grating at the side of the grey dot. The location of the coloured dot changed every four trials, while the actual colour of the dot (cyan or orange) changed pseudo-randomly (counterbalanced such that both colours occurred equally often at both locations in each block) from trial to trial. The colour of the dot predicted the overall orientation of the grating stimuli that would be presented on the attended side (45° or 135°) with 75% validity. For instance, a cyan dot presented on the left could predict with 75% validity that a pair of 45° gratings would be presented on the left side of the screen. On the remaining 25% of trials, a pair of gratings with the orthogonal (unexpected) orientation would be presented. Which colour predicted which orientation was counterbalanced across participants. The unattended (non-cued) gratings could either have the same (50%) or different orientation (50%) from the attended (cued) gratings (counterbalanced). In other words, the predictive cue contained no information about the likely orientation of the non-cued grating.

Within each pair of gratings, the second grating always differed slightly from the first in terms of orientation and contrast (see ‘Stimuli’). In separate runs (128 trials, ~14 minutes), participants performed an orientation discrimination task or a contrast discrimination task on the cued gratings. In the orientation task, participants had to judge whether the second cued grating was rotated clockwise or anti-clockwise with respect to the first cued grating (Figure 1). In the contrast task, participants had to judge whether the second cued grating had higher or lower contrast than the first cued grating. In both tasks, participants were instructed to respond within 1s of the disappearance of the second set of gratings – if they missed this deadline the fixation point briefly turned red (100ms) to indicate a missed response. The direction of rotation and contrast change for the non-cued grating were independent of that of the cued grating. The two gratings also differed in spatial phase and spatial frequency (see ‘Stimuli’), to prevent participants from using motion cues in performing the orientation task. Each participant completed a total of four runs (two of each task, the order was counterbalanced over participants) of the experiment, yielding a total of 512 trials. Each run consisted of two blocks of 64 trials, separated by a 30 second break during which the screen was blank. For each participant, the orientation and contrast differences between the two cued gratings were determined by an adaptive staircase procedure^16^ separately for the orientation and the contrast task. The staircase values were determined during the preceding behavioural session (see below) and checked during a short practice block in the MRI-scanner. During the fMRI session, these orientation and contrast differences were set as fixed values.

Participants were also exposed to a functional localiser consisting of full-field flickering gratings, in order to allow multivariate pattern analysis of orientation-specific BOLD signals. During the localiser, a fixation bull’s eye was presented, surrounded by circular (diameter = 18°) flickering gratings. These gratings were presented at 100% contrast in blocks of 15.38 s (10 TRs). The gratings were flickering on and off at a frequency of 2 Hz. Each block contained gratings with a fixed orientation of 45° or 135° and random spatial phase and spatial frequency (1.0 or 1.5 cycles/°). One cycle of the localiser consisted of four such grating blocks (either 45-135-45-135 or 135-45-135-45, randomly) and one fixation block (15.38 s), in which the screen was blank except for the fixation bull’s eye. This was repeated 8 times and lasted ~10 minutes. Concurrently, a stream of green letters was presented in the fixation bull’s eye. Participants were instructed to press a button whenever they detected either an ‘X’ or a ‘Z’ in this letter stream. This task was meant to ensure fixation and to avoid eye movements to the flickering gratings.

At the end of the experiment, we included a population receptive fields (pRFs) mapping session, in order to characterise the receptive field properties of early visual cortical areas and to allow delineation of the borders between retinotopic areas in early visual cortex^17,18^. During these runs, bars containing full contrast flickering checkerboards (2 Hz) moved across the screen in a circular aperture with a diameter of 18°. The bars moved in eight different directions (four cardinal and four diagonal directions) in 20 steps of 0.9°, one step per TR (1538 ms). Four blank fixation screens (4 TRs, 10.8 s) were inserted after each of the cardinally moving bars. Throughout each run (~5 min), a coloured fixation dot was presented in the centre of the screen, changing colour (red to green and green to red) at random time points. Participants’ task was to press a button whenever this colour change occurred. Participants performed two to four runs of this task.

### Behavioural session

Prior to the fMRI session (1-4 days), all participants completed a behavioural session. The aim of this session was to familiarise participants with the colour-grating associations and to determine the staircase values for both the orientation and the contrast discrimination task (see above).

In order to familiarise participants with the colour-grating associations, they first performed an orientation identification task. In this task, visual stimulation was identical to that in the main experiment, except that just one set of gratings (one left and one right) was presented, rather than two. Participants’ task was simply to report the orientation of the cued grating (45° or 135°), within 750 ms of grating offset. Note that in this task, as opposed to those performed in the main experiment, the cue predicted both the likely orientation of the grating and the associated response (75% valid). This task was used during the behavioural session only to promote learning of the colour-grating associations, as well as allowing us to check whether learning had occurred by comparing reaction times and accuracy to expected and unexpected grating orientations. All participants completed 3 blocks of 128 trials (~20 min in total) of this task.

After these blocks of the orientation identification task, participants were instructed about the orientation and the contrast discrimination tasks. Next, they completed a few practice trials in which they received feedback on their performance, followed by one practice block of both tasks in which they did not receive any feedback. After these practice blocks, performed four blocks of 128 trials of each task (the order was counterbalanced over participants), during which the orientation and contrast differences were determined by an adaptive staircase procedure^16^ set to an overall correct response percentage of 75%. The obtained staircase values were used during the fMRI session (see above).

### fMRI acquisition

Functional images were acquired using a 3D echo-planar imaging sequence (TR = 1538 ms, TE= 25 ms, 64 transversal slices with a distance of 50%, voxel size of 2 × 2 × 2 mm, FoV read of 256 and FoV phase of 100%, GRAPPA acceleration factor 2, 15° flip angle). Anatomical images were acquired using a T1-weigted MP-RAGE sequence, using a GRAPPA acceleration factor of 2 (TR = 2300 ms, TE = 3.03 ms, voxel size 1 × 1 × 1 mm, 192 transversal slices, 8° flip angle).

### fMRI preprocessing

The images were preprocessed using SPM8 (www.fil.ion.ucl.ac.uk/spm, Wellcome Trust Centre for Neuroimaging, London, UK). The first four volumes of each run were discarded to allow T1 equilibration. The functional images were spatially realigned to the mean image. Head motion parameters were used as nuisance regressors in the general linear model (GLM). Finally, the structural image was coregistered with the mean functional volume.

### Population receptive field (pRF) estimation and retinotopic mapping

The data from the moving bar runs were used to estimate the population receptive field (pRF) of each voxel in the functional volumes, using MrVista (http://white.stanford.edu/software/). In this analysis, a predicted BOLD signal is calculated from the known stimulus parameters and a model of the underlying neuronal population. The model of the neuronal population consisted of a two-dimensional Gaussian pRF, with parameters *x0, y0*, and *σ*, where *x0* and *y0* are the coordinates of the centre of the receptive field, and *σ* indicates its spread (standard deviation), or size. All parameters were stimulus-referred, and their units were degrees of visual angle. These parameters were adjusted to obtain the best possible fit of the predicted to the actual BOLD signal. For details of this procedure, see Dumoulin and Wandell^17^. By means of this method, we were able to estimate the coordinates of the receptive field centre, as well as the receptive field size, of each voxel. Once estimated, *x0* and *y0* were converted to eccentricity and polar-angle measures, which were overlayed on inflated cortical surfaces using Freesurfer (http://surfer.nmr.mgh.harvard.edu/) to identify the boundaries of retinotopic areas in early visual cortex^18,19^. In this way, we identified areas V1, V2 and V3 for the left and right hemisphere separately. For all analyses discussed below, only voxels for which the pRF estimation provided a good fit (i.e. variance explained > 20%) were included.

### Main task BOLD amplitude analysis

The main task data of each participant were modelled using an event-related approach, within the framework of the GLM. We distinguished 16 different conditions, according to the following four two-level factors: 1) Cued location (left or right), 2) Expected vs. Unexpected grating orientation at the cued location, 3) Cue congruent vs. Cue incongruent grating orientation at the non-cued location 4) Task. Regressors representing the different conditions were constructed by convolving delta functions with peaks at the onsets of the first grating pair in each trial with a canonical haemodynamic response function (HRF) and its temporal and dispersion derivatives^20,21^. Instruction and break screens were included as regressors of no interest, as were head motion parameters^22^. The peak of the fitted BOLD response, per condition, was converted to percent signal change by dividing by the mean signal of each run, and subsequently averaged over the different runs. (Note that we determined the peak latency based on the fitted BOLD response collapsed over all conditions and all runs, per participant.)

Within each visual area (V1, V2, and V3, per hemisphere), we defined a region of interest (ROI) consisting of voxels with a pRF centre on the grating location contralateral to the cortical hemisphere (i.e. right grating location for left hemisphere V1). BOLD amplitude, as determined by the GLM analysis described above, was averaged over the voxels within each ROI. Note that each trial yielded two data points of interest: the BOLD response to the cued grating, and the BOLD response to the non-cued grating. In order to be able to collapse across the left and right hemispheres, we defined the following four two-level factors for each grating (rather than for each trial, as in the GLM above): 1) Cued/Noncued – whether the grating is cued, i.e. attended, or not; 2) Congruent/Incongruent – whether the grating’s orientation is congruent with the expectation cue or not; 3) Contra/Ipsi – whether the grating is contra- or ipsilateral to the current ROI; 4) Task – orientation or contrast discrimination. After averaging across the left and right hemisphere ROIs for each visual area (V1, V2, and V3), the resulting BOLD amplitudes were subjected to a 4-way repeated measures ANOVA with the four factors defined above.

In order to test whether any obtained effects were specific to the grating locations or extended into non-stimulated visual cortex, we also defined a background ROI, consisting of voxels with a pRF on the background, non-overlapping with either of the grating locations (separately for V1, V2 and V3). Given that we measured pRFs using moving bars presented in a 18° diameter circular aperture, only voxels responding to stimulation within 9° of fixation had a good pRF fit and were included in the ROIs. Thus, the background ROI consisted of a 18° diameter circle around fixation, minus the two grating locations. In order to test for effects of our experimental manipulations in this ROI, we tested a 3-way repeated measures ANOVA with the same factors as above, except for Contra/Ipsi, since this ROI was collapsed over the two hemispheres. Furthermore, we tested whether effects differed significantly between the grating ROIs and the background ROI through a 4-way repeated measures ANOVA with the same factors as the 4-way ANOVA applied to the grating ROIs, except that the two-level factor ‘Contra/Ipsi’ was replaced with a three-level factor ‘Cortical location’, the levels being: 1) grating ROI contralateral to grating, 2) grating ROI ipsilateral to grating, 3) background ROI. Finally, we performed an ANOVA with factors ‘Cued/Non-cued’, ‘Congruent/Incongruent’, ‘Contra/Ipsi’ and ‘Visual area’, to investigate whether effects differed between the visual areas (V1, V2 and V3). All statistical analyses were based on these ROI data. Error bars in all figures indicate within-subject standard error of the mean^23,24^ (SEM).

### Orientation specific BOLD signal analysis

In order to inspect orientation specific BOLD signals, we applied the same GLM as described above, but creating two regressors for every condition, one for each grating orientation (45° or 135°). In order to establish voxel’ orientation preference, we also applied a GLM analysis to the functional localiser, with one regressor for 45° gratings and one for 135° gratings. Within each ROI as defined above, we selected the 25% of voxels with the greatest preference for 45° and the 25% of voxels with the greatest preference for 135°. We then subtracted the BOLD response in the 135° preferring voxels from the response in the 45° voxels and used this in further analyses as our orientation specific BOLD signal. For each condition, we subsequently subtracted the orientation specific BOLD signal for the trials in which a 135° grating was presented from the BOLD signal for the trials in which a 45° grating was presented. As a consequence of these subtractions, we expect a significantly positive value for conditions in which an orientation specific BOLD signal is present, and a value of zero when no such signal is present. One advantage of performing this double subtraction is that we end up with the exact same experimental factors as for the BOLD amplitude analysis described above, but now with orientation specific BOLD signal (as opposed to overall BOLD amplitude) as the dependent variable. Hence, all statistical analyses on the orientation specific BOLD signal were performed identically (i.e., using a 4-way repeated measures ANOVA) to the BOLD amplitude analysis described above.

### Retinotopic reconstruction of BOLD response

In addition to ROI specification, the estimated pRF parameters allowed a straightforward and intuitive reconstruction of the BOLD response from cortical space to visual space^25,26^ (Figure 1C). Each voxel’s receptive field can be represented by a two dimensional Gaussian, with peak coordinates (*x0, y0*) and standard deviation *σ*. The reconstruction in visual space consisted of the sum of the 2D-Gaussians of all voxels in a given visual area, weighted by the voxels’ BOLD response:

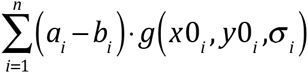

Where *n* is the number of voxels in a given area, *a_i_*, and *b_i_*, are responses to certain conditions, both in percent signal change, and *g*(*x0_i_,y0_i_,σ_i_*) is the 2D-Gaussian defining the voxel’s receptive field. The rationale is the following: if a voxel in V1 is highly active, then this reflects activity in V1 neurons corresponding to the region of visual space modelled by the 2D-Gaussian. Therefore, in order to represent this activity in visual space, we multiplied the voxel’s 2D-Gaussian receptive field with its activity (i.e., BOLD signal) and projected the result on a 2D map of visual space. By doing this for all V1 voxels, we obtained a reconstruction of the BOLD signal in V1 in visual space. We also calculated a ‘baseline’ map of the visual field, with each voxel’s weight set to 1:

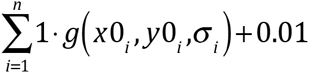

We divided all reconstructions by this baseline map, pixel wise, in order to compensate for the relatively higher number of voxels near the fovea than in the periphery (i.e., cortical magnification). Finally, all reconstructions were spatially smoothed with a 2-D Gaussian smoothing kernel with *σ* = 0.5°. These reconstructions were used to visualise the BOLD response, while statistical analyses were performed on the ROI data described above (‘Main task BOLD amplitude analysis’).

## Results

### Behavioural results

In order to familiarise participants with the predictive relationship between the coloured cues and the orientation of the grating stimuli, participants performed an orientation identification task during the behavioural session prior to the fMRI session. In this task, participants simply reported the orientation of the cued grating (45° or 135°), which would be preceded by either a valid (75%) or invalid predictive cue (25%; see Materials and Methods for details). Participants were more accurate and faster to identify the orientation of expected (i.e. validly cued) gratings than unexpected gratings (mean accuracy = 89.4% vs. 84.3%, *t*_22_ = 3.53, *p* = 0.002; mean RT = 451 ms vs. 464 ms, *t*_22_ = -2.46, *p* = .022), demonstrating that participants learned the colour-orientation associations correctly. During the fMRI session, participants performed an orientation discrimination task and a contrast discrimination task, in separate blocks (see Materials and Methods). Participants were able to discriminate small differences in orientation (4.0° ± 2.0°, accuracy = 83.3% ± 7.4%, mean ± SD) and contrast (7.6% ± 2.3%, accuracy = 77.6% ± 9.5%, mean ± SD) of the cued gratings. Accuracy was significantly higher for the orientation task than for the contrast task (F_1,22_ = 4.94, *p* = 0.037), but there was no difference in reaction time between the tasks (mean RT = 702 ms vs. 711 ms, *F*_1,22_ = 0.30, *p* = .59). Overall, accuracy and reaction times were not influenced by whether the cued grating had the expected or the unexpected orientation (accuracy: *F*_1,22_ = 0.0014, *p* = 0.97; RT: *F*_1,22_ = 0.066, *p* = 0.80), nor was there an interaction between task (orientation task vs. contrast task) and expectation (expected vs. unexpected) (accuracy: *F*_1,22_ = 0.64, *p* = 0.43; RT: *F*_1,22_ = 0.65, *p* = 0.43). Note that, unlike for the orientation identification task during the behavioural session, these discrimination tasks were orthogonal to the expectation manipulation, in the sense that the expectation cue provided no information about the likely correct choice.

### BOLD amplitude results

Below, we present the spatially specific responses to the grating stimuli in early visual cortex in two different ways. First, we used a reconstruction method to visualise the BOLD response to the grating stimuli in retinotopic space (Figure 2). Second, we employed a more conventional ROI approach, selecting groups of voxels based on the location of their receptive field and averaging their BOLD response (Figure 3). These analysis strategies are complementary: while the first method allows for a characterization and visualisation of neural activity concurrently for all parts of visual space, the second approach allows for a more straightforward statistical quantification of the experimental effects.

**Figure 2.**
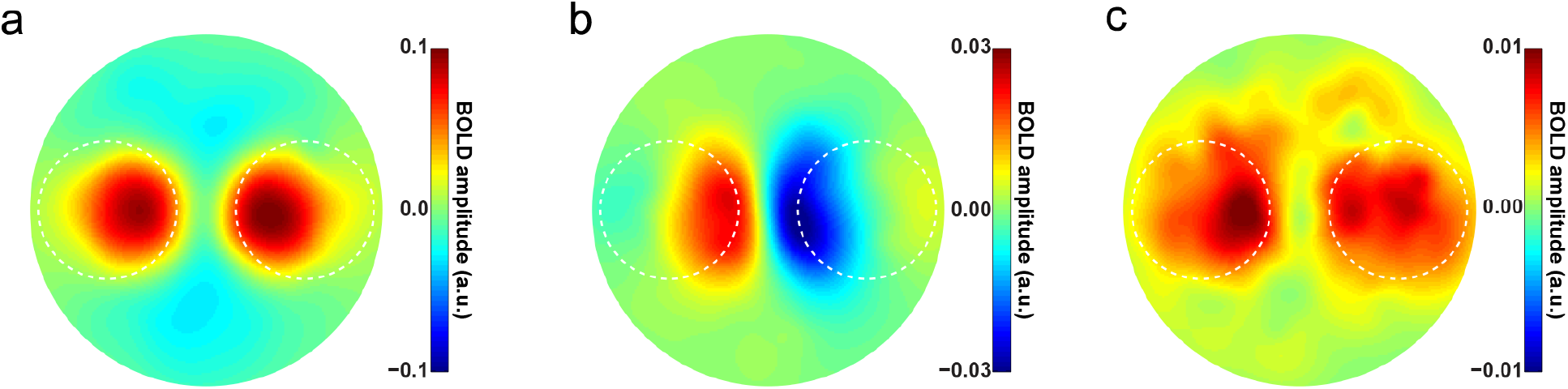
Retinotopic reconstructions of attention and expectation effects. **(a)** Reconstruction of the BOLD response to the grating stimuli in V1, based on the contrast ‘All trials – baseline’. Dashed white circled indicate the location of the gratings, in retinotopic space. **(b)** Reconstruction of the spatial attention effect, based on the contrast ‘Attention left – Attention right’. **(d)** Reconstruction of the expectation effect. That is, ‘Cued grating Unexpected – Expected’. We collapsed over trials in which the left and right gratings were cued by first flipping the reconstructions of the ‘cued right’ trials horizontally, to move them into the reference frame of the ‘cued left’ trials, before averaging. See Supplementary Figure S1 for separate reconstructions for ‘cued left’ and ‘cued right’ trials.

**Figure 3.**
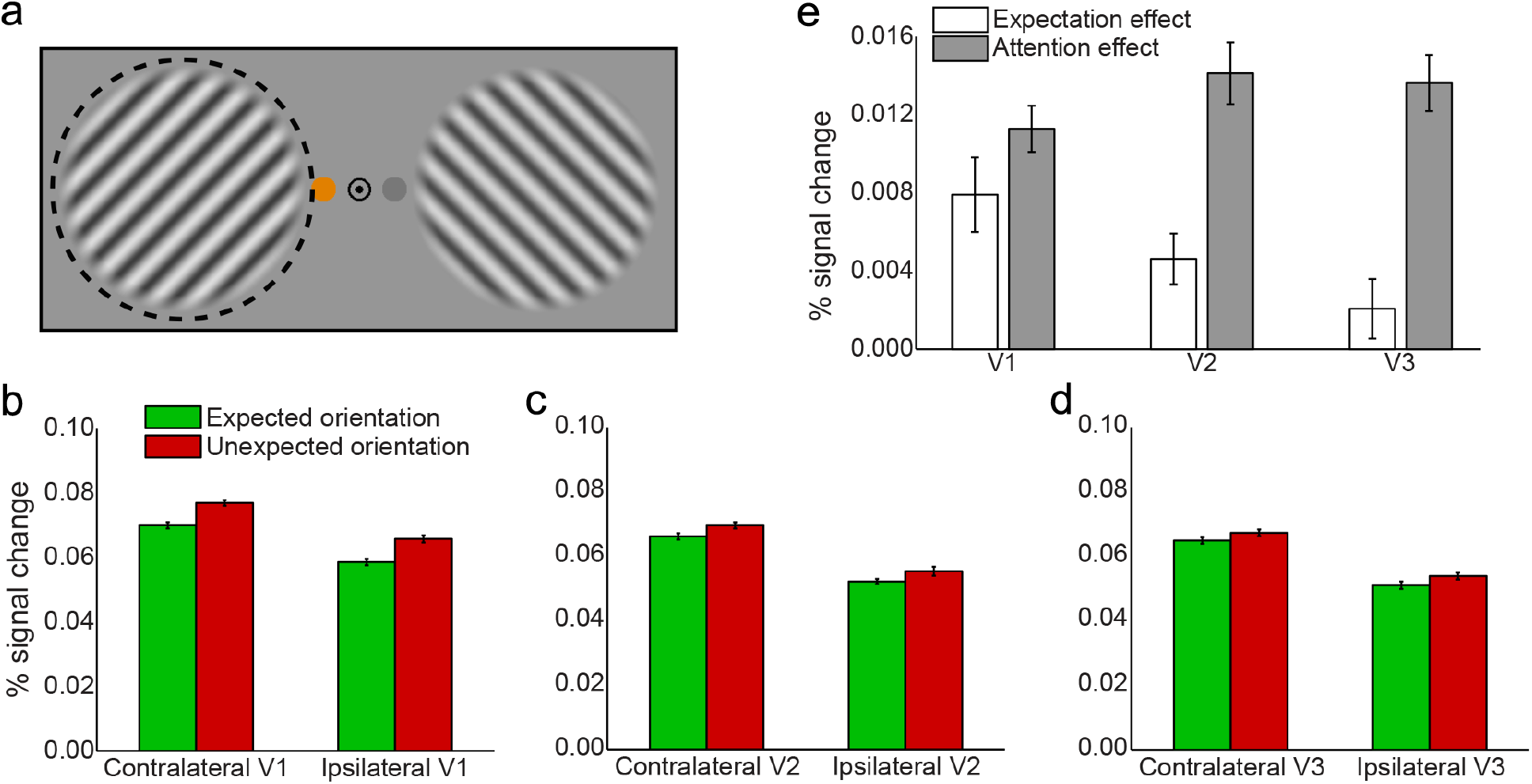
BOLD amplitude evoked by expected and unexpected gratings. **(a)** Here, trials are split up according to whether the gratings on the cued side had the expected or unexpected orientation. BOLD amplitude is shown separately for the cortical ROI in which the grating was processed (i.e. contralateral hemisphere) as well as for the ROI in the opposite (i.e. ipsilateral) hemisphere. Results are shown separately for **(b)** V1, **(c)** V2, and **(d)** V3. Note that the BOLD response is higher contralateral to the cued grating than contralateral to the non-cued grating (i.e., ipsilateral to the cued grating), as a result of spatial attention. Further, the BOLD response is lower when the cued grating has the expected orientation (green bars) than when it has the unexpected orientation (red bars). **(e)** Effects of expectation (white bars) and spatial attention (gray bars) per visual region. Error bars indicate within-subject SEM. (See Supplementary Figure S3 for BOLD amplitude conditioned on whether the non-cued grating was congruent or incongruent with the expectation cue.)

The location of the spatial attention cue had a strong effect on the BOLD response evoked by the gratings in V1 (Figure 3B; *F*_1,22_ = 90.93, *p* = 2.8 × 10^-9^). Notably, this attention modulation was strongest peri-foveally, and weakest in voxels representing the most peripheral part of the grating (compare Figure 2B to Figure 2A).

Additionally, the BOLD response evoked by the gratings in V1 was strongly modulated by expectation; when the orientation of the cued grating was unexpected, this led to a higher BOLD response than when the orientation of the cued grating matched the expectation (Figure 3B; *F*_1,22_ = 17.19, *p* = 4.2 × 10^-4^). This effect had a striking bilateral distribution (Figure 2C); the increased BOLD response was not specific to the cortical location in which the cued grating was processed (*F*_1,22_ = 0.62, *p* = 0.44), but was present at both the cortical locations processing the cued and the non-cued gratings (Figure 2C and Supplementary Figure S1). However, the effect did not spread to the non-stimulated background (*F*_1,22_ = 0.37, *p* = 0.55; Supplementary Figure S2), but was specific to the cortical locations processing the gratings (Figure 2C; *F*_2,44_ = 15.18, *p* = 9.7 × 10^-6^). In other words, when the grating at the cued side violated the orientation expectation, this led to an increased BOLD signal at *both* grating locations, but not elsewhere in the visual field.

Importantly, the expectation effect was only contingent on the orientation of the *cued* grating (F_1,22_ = 17.19, *p* = 4.2 × 10^-4^); whether or not the orientation of the *non-cued* grating (i.e. the unattended grating, whose orientation was independent of the expectation cue) was congruent or incongruent with the expectation cue had no effect on the BOLD response (Supplementary Figure S3; *t*_22_ = 0.59, *p* = 0.56). In other words, the expectation effect was spatially specific in the sense that only the orientation at the cued location had an impact on the BOLD response, but spatially non-specific in the sense that the increased BOLD response to an expectation violation was present at both gratings’ cortical locations.

In addition to V1, spatial attention also strongly modulated the BOLD response in V2 (*F*_1,22_ = 79.56, *p* = 9.3 × 10^-9^) and V3 (*F*_1,22_ = 89.79, *p* = 3.2 × 10^-9^), while expectation significantly affected V2 (*F*_1,22_ = 12.66, *p* = 0.0018) but not V3 (*F*_1,22_ = 1.85, *p* = 0.19). In fact, the effects of spatial attention and expectation seemed to behave differently when moving up the visual cortical hierarchy (Figure 3E): The effect of expectation diminished significantly from V1 to V3 (*F*_2,44_ = 11.11, *p* = 1.2 × 10^-4^), being stronger in V1 (Figure 3B) than in V2 (Figure 3C; t_22_ = 2.58, *p* = 0.017) and V3 (Figure 3D; *t*_22_ = 4.08, *p* = 5.0 × 10^-4^), and stronger in V2 than in V3 (*t*_22_ = 2.62, *p* = 0.016), while the effect of spatial attention, on the other hand, significantly *increased* going up the cortical hierarchy (*F*_2,44_ = 3.99, *p* = 0.026), being smaller in V1 than in V2 (*t*_22_ = 2.37, *p* = 0.027) and V3 (*t*_22_ = 2.21, *p* = 0.038). This latter increase is in line with previous studies showing effects of attention being stronger in higher-order visual cortex than in lower-order areas^27,28^. In sum, the effects of expectation and spatial attention behaved differently across the cortical hierarchy, with expectation effects being strongest, and attention effect being weakest, in V1 (Figure 3E).

The type of task participants performed (orientation or contrast discrimination) had no effect on the BOLD response, nor did it interact with any of the other factors (i.e. attention or expectation) (all *p* > 0.10). This is in line with results from previous studies employing these tasks^3,11^, and suggests that the effects of orientation expectation are independent of the task-relevance of orientation.

### Orientation specific BOLD results

The orientation specific analysis revealed a significant orientation specific BOLD response to gratings in the contralateral hemisphere in V1 (*F*_1,22_ = 14.77, p = 8.8 × 10^-4^), but not in V2 (*F*_1,22_ = 1.59, *p* = 0.22) and V3 (*F*_1,22_ = 0.82, *p* = 0.37). In V3, the orientation signal was higher for gratings that matched than those that mismatched the expectation cue (*F*_1,22_ = 6.20, *p* = 0.021), but no such effect was present in V1 (*F*_1,22_ = 4.1 × 10^-4^, *p* = 0.98) and V2 (*F*_1,22_ = 3.51, *p* = 0.074). In V2, attended gratings evoked a larger orientation signal in the contralateral hemisphere than unattended gratings (*F*_1,22_ = 5.03, *p* = 0.035), but this effect was absent in V1 (*F*_1,22_ = 0.62, *p* = 0.44) and V3 (*F*_1,22_ = 0.074, *p* = 0.79). However, neither the effect of expectation (*F*_2,44_ = 2.58, *p* = 0.087) or spatial attention (*F*_2,44_ = 1.24, *p* = 0.30) significantly differed between regions, suggesting that the inconsistencies between regions may not be reliable, and more likely due to noise. In fact, when collapsing across regions, potentially increasing the signal-to-noise ratio, there was a significant increase in orientation signal for gratings that matched, compared to those that mismatched, the expectation cue (*F*_1,22_ = 4.98, *p* = 0.036). There was no significant effect of spatial attention (*F*_1,22_ = 2.67, *p* = 0.12) across regions. Taken together, the inconsistency of these effects suggests that the orientation specific BOLD signal had a relatively low signal-to-noise ratio, precluding strong conclusions on the basis of these analyses. Finally, there was no significant orientation specific BOLD signal in the unstimulated background, nor were there any effects of attention or expectation (all *p* > 0.05). Similarly, applying a different multivariate analysis method, linear support vector machines (SVM), did not yield any significant effects of attention and expectation in any of the ROIs.

## Discussion

Prior expectations can be specific to both *what* you will see and *where* you will see it. For instance, when you open your front door you not only know what colour sofa to expect, but also where it is located. Here, we found that when the orientation of a visual stimulus is different than expected, this leads to an increased neural response in early visual cortex both in cortical locations processing the unexpected stimulus, as well as in cortical locations processing an independent and irrelevant stimulus. Importantly, this increased neural response was evoked only when the stimulus to which the expectation cue pertained had an unexpected feature; whether or not the features of an independent, non-cued stimulus were congruent with the expectation cue did not affect the BOLD response. In other words, the effect of feature-based expectation was spatially specific in the sense that it was only evoked by the stimulus that the expectation cue pertained to. If feature-based expectation would have been spatially non-specific (as has been observed for feature-based attention^9–11^) neural responses should also have differed for the non-cued stimulus, depending on whether its orientation matched or mismatched the expectation of the cued stimulus. This is not in line with our findings, suggesting that expectations are both spatially and feature specific^29^.

The *consequences* of a violated expectation, however, were spatially non-specific, in that it evoked an increased neural response in both cortical hemifields processing a stimulus (but not in the non-stimulated background). This is a surprising finding that is not readily explained by current theories of feature-based expectation or attention. What might be the neural mechanism underlying this bilateral neural effect? One possibility is that a violation of the expectation by the first grating at the cued location leads to an upregulation of visual processing across the visual field. This upregulation may take the form of a spatially non-specific gain modulation of neural responses in early visual cortex, leading to increased neural responses at all locations where a stimulus is presented. The effect of such a gain modulation triggered by the unexpected grating would be particularly visible in the current paradigm, since the first set of gratings was followed by a second set of gratings, which would (also) be affected by the non-specific gain modulation. Future studies could follow up on this by employing a paradigm with only one set of gratings per trial. Such studies could also investigate whether it is important that the independent, irrelevant stimulus is similar to the expected stimulus (as is the case here; both stimuli were oriented gratings), or whether an expectation violation also leads to modulation of the response to stimuli with completely different features (e.g., geometrical shapes or moving dots, or even auditory stimuli).

Our retinotopic reconstruction technique also revealed a particular distribution of the effect of spatial attention (Figure 2B), with effects being much stronger near the fovea than further in the periphery. This could be caused by an inhomogeneity in the attentional field, with strongest attentional gains in the parts of the grating stimuli closer to the fovea, where visual acuity is higher and therefore the largest signal benefit can be obtained.

### Expectation and attention across the visual cortical hierarchy

Apart from having different spatial distributions (cf. Figures 2B and 2C), the effects of spatial attention and feature-based expectation were also distributed over the cortical hierarchy differently. In line with previous literature, the effect of spatial attention was stronger in higher-order visual areas (V2 and V3) than in V1^27,28^. The effect of expectation, on the other hand, was strongest in V1 and significantly decreased going up the cortical hierarchy (i.e. to V2 and V3). One potential explanation for this is that effects of feature-based expectation are strongest in those cortical regions that code for the expected feature. In the case of high spatial frequency gratings, as used in the current study, the cortical region most optimised for coding the expected features is likely to be V1. In line with this, previous studies using similar stimuli have also observed modulations of activity by orientation expectation in V1, but less so in V2 and V3^1,3^. On the other hand, studies manipulating expectations of higher level stimuli, such as faces and houses^30–32^ and letter shapes^33,34^ have generally reported effects in high-level visual areas specialised in processing such stimuli. As a notable exception, studies on the top-down filling-in of missing portions of high-level scene stimuli have found strongest filling-in effects in V1^35,36^, leading to suggestions that V1 may have a special role in processing top-down expectations, as a type of high-resolution dynamic blackboard^37–39^.

### Limitations of the current study

In the current study, the cue informed both the task-relevant location as well as the expected orientation at that location. Therefore, the expectation was always about the task-relevant grating. Future studies could investigate whether similar expectation effects (i.e., a BOLD increase at both stimulus locations) would also occur when the stimulus to which the expectation pertained was task-irrelevant and unattended. Moreover, in addition to the overall amplitude of the neural response, it would be of interest to study the effects of expectation on stimulus representations across the visual field, using multivariate pattern analyses. Such analyses have been revealing in previous studies, reporting for instance opposing effects of expectation on BOLD amplitude and stimulus representations^3^, as well as demonstrating the spatial non-specificity of feature-based attention in humans^10,11^. Unfortunately, the results of the pattern analyses in the current study were inconclusive, potentially due to low signal-to-noise ratio. The signal-to-noise ratio of orientation-specific BOLD signals could be improved by 1) presenting grating stimuli in an annulus around fixation, maximising the sampling of the orientation biases in visual cortex^40,41^, and/or 2) employing a blocked rather than an event-related design^10,11^. For instance, one could study the orientation-specific BOLD signal evoked by a large grating annulus around fixation, while manipulating expectations about gratings in the periphery. This could be a fruitful approach for follow-up studies.

### Conclusions

Here, we show that when participants are presented with a stimulus that has an unexpected orientation, this leads to an increased neural response both in the cortical location processing that stimulus, as well as in another cortical location processing a similar but irrelevant stimulus. This is in line with an expectation violation leading to a spatially non-specific gain increase, upregulating the neural response to stimuli across the visual field. This effect is only evoked by the stimulus that the expectation pertains to; whether or not an independent stimulus is congruent or incongruent with a participant’s expectation about another stimulus has no effect on the BOLD response. Together, these results suggest that expectations are simultaneously feature-specific and spatially specific^29^, but a violation of the expectation can have neural consequences that are not spatially specific, possibly reflecting a gain modulation across the visual field.

## Acknowledgements

This work was supported by The Netherlands Organisation for Scientific Research (NWO) to P.K. (Rubicon grant 446-15-004) and F.d.L. (VIDI grant 452-13-016) and the James S McDonnell Foundation to F.d.L. (Understanding Human Cognition 220020373).

## Author contributions

P.K., L.L.F.v.L and F.d.L. designed the experiment, P.K. and L.L.F.v.L collected the data, P.K. and L.L.F.v.L analysed the data, P.K., L.L.F.v.L and F.d.L. wrote the paper.

## Additional information

Conflict of Interest: The authors declare no competing financial interests.

